# Structural characterization of the N-terminal domain of the *Dictyostelium discoideum* mitochondrial calcium uniporter

**DOI:** 10.1101/848002

**Authors:** Yuan Yuan, Chan Cao, Maorong Wen, Min Li, Ying Dong, Lijie Wu, Jian Wu, Tanxing Cui, Dianfan Li, James J. Chou, Bo OuYang

**Affiliations:** State Key Laboratory of Molecular Biology, CAS Center for Excellence in Molecular Cell Science, Shanghai Institute of Biochemistry and Cell Biology, Chinese Academy of Sciences; University of Chinese Academy of Sciences, 333 Haike Road, Shanghai 201203, P. R. China; Department of Biological Chemistry and Molecular Pharmacology, Harvard Medical School, Boston, MA 02115, USA; Shanghai Institute for Advanced Immunochemical Studies and iHuman Institute, ShanghaiTech University, Shanghai 201210, China; Ninth People’s Hospital, Shanghai Jiao Tong University School of Medicine, Shanghai, 200125, China

**Keywords:** mitochondrial calcium uniporter (MCU), *Dictyostelium discoideum*, N-terminal domain, crystal structure, oligomerization

## Abstract

The mitochondrial calcium uniporter (MCU) plays a critical role in the mitochondrial calcium uptake into the matrix. In metazoans, the uniporter is a tightly regulated multi-component system including the pore-forming subunit MCU and several regulators (MICU1, MICU2, EMRE). The calcium-conducting activity of metazoan MCU requires the single-transmembrane protein EMRE. *Dictyostelium discoideum* (Dd), however, developed a simplified uniporter for which the pore-forming MCU (DdMCU) alone is necessary and sufficient for calcium influx. Here, we report a crystal structure of the N-terminal domain (NTD) of DdMCU at 1.7 Å resolution. The DdMCU-NTD contains four helices and two strands arranged in a fold that is completely different from the known structures of other MCU-NTD homologs. Biochemical and biophysical analyses of DdMCU-NTD in solution indicated that the domain exists as oligomers, most probably as a pentamer or hexamer. Mutagenesis showed that the acidic residues Asp60, Glu72 and Glu74, which appeared to mediate the parallel interface as observed in the crystal structure, participated in the self-assembly of DdMCU-NTD. Intriguingly, the oligomeric complex readily dissociated to lower-order oligomers in the presence of calcium. We propose that the calcium-triggered dissociation of NTD regulates the channel activity of DdMCU by a yet unknown mechanism.

## Introduction

Mitochondrial calcium (Ca^2+^) homeostasis plays a vital role in regulating various cellular activities, such as aerobic metabolism and cell apoptosis (Denton and McCormack, 1980). Several mitochondrial calcium transporters have been identified to be responsible for the maintenance of an internal steady state of calcium ions within the cytoplasm (Cai and Lytton, 2004; Jiang et al., 2009; Palty et al., 2004). The Ca^2+^ uniporter activity responsible for mitochondrial Ca^2+^ uptake was reported more than 50 years ago (Deluca and Engstrom, 1961; Vasington and Murphy, 1962) and later this uniporter was demonstrated by electrophysiological measurements to be an ion channel with remarkably high conductance and selectivity for calcium (Kirichok et al., 2004). It was not until eight years ago that the molecular components of the uniporter had been identified (Baughman et al., 2011; De Stefani et al., 2011; Perocchi et al., 2010; Raffaello et al., 2013; Sancak et al., 2013). In vertebrates, the uniporter is a holocomplex consisting of multiple subunits, including two membrane-spanning subunits and two soluble proteins in the intermembrane space. The two soluble proteins are the mitochondrial calcium uptake proteins, MICU1 and MICU2, which are Ca^2+^ sensing proteins that gate the activity of the pore in response to Ca^2+^concentrations outside the mitochondria (Ahuja and Muallem, 2014; Kamer and Mootha, 2014, 2015a;Patron et al., 2014; Petrungaro et al., 2015). The two membrane proteins are the mitochondrial calcium uniporter (MCU) that forms the Ca^2+^ conducting channel and the small single-pass TM protein EMRE (<u>E</u>ssential <u>M</u>CU <u>RE</u>gulator) (Sancak et al., 2013; Vais et al., 2016). As its name suggests, EMRE is essential for activating or keeping the pore in an open conformation, as well as for transducing the Ca^2+^sensing from MICU1/2 to MCU (Sancak et al., 2013). Now that the key molecular components of the calcium uniporter have been identified, more attentions are turning to the molecular mechanism of MCU function and regulation (Kamer and Mootha, 2015b).

The molecular architecture of the MCU core domain was first revealed by solution NMR and negative stain electron microscopy (EM) studies, with a truncated *C. elegans* (Ce) MCU protein that lacks the extramembrane N-terminal domain (NTD; residues 1-165) (Oxenoid et al., 2016). The protein, named CeMCU-ΔNTD, contains two transmembrane (TM) helices and adapts a pentameric complex with the second TM helix forming a hydrophilic pore across the membrane. The DxxE sequence motif critical for channel activity (De Stefani et al., 2015) between the first and the second TM helices forms two carboxylate rings at the entry of the pore, featuring a Ca^2+^ selectivity filter (Cao et al., 2017). In the NMR study the NTD was not included, a separate X-ray study determined the high resolution structure of the NTD of the human MCU (HsMCU), which adopts a ubiquitin-like fold (Lee et al., 2015). Another crystal structure of HsMCU_72-189_ also revealed a β-grasp-like fold with a negatively charged region that interacts with divalent cations and this acidic patch is important for regulating human MCU activity in response to Mg^2+^ and Ca^2+^ binding (Lee et al., 2016). Extensive efforts have been put to obtain the structures of the full-length MCU since its discovery. In 2018, the overall structures of the MCU were independently solved by several groups within a couple of months. The full-length structures of the fungal MCU homologues of *Fusarium graminearum* (Fg) and *Metarhizium acridum* (Ma), have been determined using X-ray crystallography and cryo-electron microscopy (cryo-EM), revealing a dimer of dimer assembly of MCU (Fan et al., 2018). Meanwhile, the structures of the full-length MCU homologues from *Neurospora crassa* (Nc), zebrafish, *Cyphellophora europaea* (Cy), and *Neosartorya fischeri* (Nf) have also been determined using cryo-EM, showing a similar tetrameric fold (Baradaran et al., 2018; Nguyen et al., 2018; Yoo et al., 2018). These homologues show high sequence similarity in their transmembrane domain and share a similar pore architecture for calcium conductivity. Remarkably, the NTDs of these homologues also share high structural similarity despite of the low sequence homology between NTDs. Recently, Wang et al. reported the cryo-EM structure of the human MCU-EMRE complex, providing the structural features for the EMRE regulation on the conformation of HsMCU (Wang et al., 2019). It is interesting to note that the NTD adopts a unique side-by-side configuration to mediate the dimerization of two HsMCU tetramers, which indicates that the NTD may play an important role in modulating the function of MCU, although the removal of the NTD does not affect mitochondrial Ca^2+^ uptake (Lee et al., 2015).

Here, we determined the crystal structure of the NTD of *Dictyostelium discoideum* (DdMCU) (residues 29-126) at 1.7 Å resolution. *Dictyostelium* harbors a simplified uniporter complex, consisting of only the MCU protein (DdMCU) to conduct Ca^2+^ (Sancak et al., 2013) (Fig. S1), no EMRE is required for the channel activity. It has been shown that expression of DdMCU alone is able to reconstitute uniporter activity in yeast mitochondria (Kovacs-Bogdan et al., 2014). The same study showed that DdMCU could rescue the mitochondrial Ca^2+^ uptake defects in HsMCU or HsEMRE knockout HEK-293T cells. Unexpectedly, the crystal structure of DdMCU-NTD showed a helix-rich fold that is completely different from the β-grasp-like structures of the HsMCU-NTD and other homologues. The structure also revealed a noncrystallographic interface including intermolecular interactions of helix α3 and loop of monomer-A with helix α2 of monomer-B. We further showed that the DdMCU-NTD formed an oligomer in solution, most probably a pentamer or hexamer. Mutating residues within the noncrystallographic interface caused the dissociation of the oligomer. Furthermore, the oligomeric state could be disrupted by the addition of Ca^2+^. Our results revealed the divergent structures of MCU NTDs while suggesting the key residues of NTD in DdMCU channel assembly.

## Materials and methods

### Protein expression and purification

The *Dictyostelium discoideum* MCU N-terminal domain (DdMCU-NTD) was cloned into the vector pET21a with a hexahistidine (6×His) at the N-terminus. The plasmid was then transformed into *Escherichia coli* BL21 (DE3) and cultured in Luria-Bertani (LB) medium at 37 °C until the optical density at 600 nm (OD_600_) reached 0.7. The expression of DdMCU-NTD was induced by the addition of 0.15 mM IPTG, followed by incubation at 18 °C for 12 hr.

The bacteria were harvested by centrifugation and resuspended in lysis buffer (200 mM NaCl, 0.2 mM PMSF, 50 mM Tris-HCl pH 8.0). After sonication, the lysate was centrifuged at 39,000 g for 40 min. Then the supernatant was passed through a Ni-NTA affinity column, washed and eluted with elution buffer (200 mM NaCl, 0.2 mM PMSF, 400 mM imidazole, 50 mM Tris-HCl pH 8.0). The imidazole was removed by dialyzing the sample against 15 mM NaCl, 5 mM DTT, and 20 mM Tris-HCl pH 8.5. The protein was then applied to a HiTrap-Q HP column (GE Healthcare) equilibrated with 15 mM NaCl, 5 mM DTT, 20 mM Tris-HCl pH 8.5 and eluted with a linear gradient to 1 M NaCl in 30 minutes. The protein was further purified using a HiLoad16/60 Superdex200 column (GE Healthcare) equilibrated with size-exclusion chromatography buffer (75 mM NaCl, 5 mM DTT, 20 mM HEPES pH 7.5). The pooled fractions were concentrated to 20 mg/mL for crystallization. All the mutants were purified following the same protocol.

For crystal structure determination, selenomethionyl (SeMet) DdMCU-NTD was expressed in *E. coli* BL21 (DE3) cells grown in M9 medium supplemented with SeMet, as described previously (Walden, 2010) The SeMet labeled mutant (L8M/L20M) was then produced by following the same purification protocol used for the wild-type protein.

### Crystallization and data collection

The DdMCU-NTD protein (20 mg/mL in 75 mM NaCl, 5 mM DTT, 20 mM HEPES pH 7.5) was crystallized using the sitting drop vapor diffusion method with a reservoir solution of 20 % (w/v) PEG 4000, 10 % (v/v) 2-propanol, 10 mM HEPES pH 7.5. Ca^2+^ was not added to the crystallization buffer and no EDTA was added either to sequester residual Ca^2+^ constitutively present in water. Crystals were grown for 3-5 weeks at 4°C by mixing equal volumes (0.5 μL) of protein and reservoir solution in MRC 2 drop 96 well crystallization plate (Swissci, Neuheim, Switzerland) and cryocooled in a cryoprotectant consisting of the reservoir solution supplemented with 35% (v/v) ethylene glycol. The SeMet crystals were produced in the same conditions used for the wild-type protein, with the exception that 8 mg/mL SeMet protein was used for crystallization. The SeMet crystals were flash-frozen in liquid nitrogen with the same cryoprotective reagents. X-ray diffraction data were collected at the BL18U beamline in the Shanghai Synchrotron Radiation Facility (SSRF). Diffraction data were processed using the XDS package (Kabsch, 2010a, b, 2014). Data collection statistics are shown in Table 1.

**Table 1.** Data collection, phasing and refinement statistics

|  | **DdMCU-WT** | **DdMCU-SeMet** |
| --- | --- | --- |
| *Data collection* |  |  |
| Space group | P 2<sub>1</sub> 2<sub>1</sub> 2<sub>1</sub> | P 1 2<sub>1</sub> 1 |
| *Cell dimensions* |  |  |
| *a*, *b*, *c* (Å) | 42.16, 59.36, 70.86 | 42.46, 70.83, 60.00 |
| $\alpha$, $\beta$, $\gamma$ (°) | 90, 90, 90 | 90, 92.17, 90 |
| Wavelength (Å) | 0.97776 | 0.97776 |
| Resolution (Å) | 45.50 - 1.68<br>(1.71 - 1.68) | 45.77 - 2.14<br>(2.20 - 2.14) |
| *R*<sub>merge</sub> | 0.050 (1.264) | 0.099 (0.823) |
| *R*<sub>pim</sub> | 0.016 (0.374) | 0.034 (0.305) |
| *I*/σ*I* | 25.5 (2.2) | 13.9 (2.8) |
| Completeness (%) | 99.7 (99.7) | 98.7 (88.6) |
| Multiplicity | 11.8 (12.3) | 9.4 (8.0) |
| *CC*<sup>*</sup> | 0.999 (0.726) | 0.998 (0.723) |
| *Refinement* |  |  |
| Resolution (Å) | 45.50 - 1.68 | 45.77 - 2.14 |
| No. reflections | 39046 | 19356 |
| *R*<sub>work</sub> / *R*<sub>free</sub> | 0.1902/0.2237 | 0.1922/0.2313 |
| No. atoms | 1595 | 3034 |
| Protein | 1477 | 2971 |
| Water | 118 | 63 |
| No. residues | 182 | 371 |
| B-factors (Å<sup>2</sup>) | 38.28 | 52.30 |
| Protein | 37.93 | 52.37 |
| Water | 42.54 | 49.01 |
| *R.m.s deviations* |  |  |
| Bond lengths (Å) | 0.008 | 0.006 |
| Bond angles (°) | 0.68 | 0.80 |
| *Ramachandran* |  |  |
| Favored (%) | 100 | 99.45 |
| Allowed (%) | 0 | 0.55 |
| Outlier (%) | 0 | 0 |
| PDB ID | 5Z2H | 5Z2I |

### Structure determination and refinement

The structure was solved using Phenix.autosol with the single-wavelength anomalous dispersion data collected using the SeMet-labeled L8M/L20M mutant. An initial model was built using Phenix.autobild (Adams et al., 2002). The structure was completed using alternate cycles of manual fitting in Coot (Emsley and Cowtan, 2004) followed by structure refinement with Phenix.refine. Detailed data collection and refinement statistics are summarized in Table 1.

### SEC-MALS analysis of the DdMCU-NTD molecular mass

The instrument setup used for the SEC-MALS experiment consisted of an Agilent 1260 Infinity Isocratic Liquid Chromatography System connected in tandem with a Wyatt Dawn Heleos II Multi-Angle Light Scattering (MALS) detector (Wyatt Technology) and a Wyatt Optilab T-rEX Refractive Index Detector (Wyatt Technology). Analytical size-exclusion chromatography was performed at room temperature using a Superdex 200 10/300 GL column (GE Healthcare) equilibrated with a mobile phase containing 75 mM NaCl, 0.3 mM NaN3, 50 mM MES pH 6.4. 100 μL purified DdMCU-NTD sample at 10 mg/mL was injected into the column and eluted at a flow rate of 0.4mL/min. The column effluent was monitored in-line with three detectors that simultaneously monitored UV absorption, light scattering and refractive index. The data from the three detectors were imported by the ASTRA software package, and the three-detector method (Slotboom et al., 2008) was used to determine the molecular mass.

### Dynamic Light Scattering

Dynamic light scattering (DLS) measurements were performed at 25 °C for DdMCU-NTD and HsMCU-NTD on a DynaPro NanoStar instrument (Wyatt) with water or buffer alone as controls. Light scattering data collection and analysis for the size and size distribution of the proteins were calculated using the Dynamics V6 software.

### Characterization of the DdMCU-NTD oligomeric state

The oligomeric state of DdMCU-NTD was first examined by chemical crosslinking using glutaraldehyde (sigma). The reaction volume of 50 μl consisted of 0.5 mg/ml protein, 25 mM HEPES-Na, pH 7.5, 150 mM NaCl and various concentrations of glutaraldehyde (0.0015%, 0.015% and 0.15%, v/v). The reaction was stopped after 2 min by adding 100 mM Tris, pH 8.0. The samples were incubated for another 5 min. The reaction mixtures were analysed using SDS-PAGE gel. Blue native gradient gel was generated with a 4-13% (w/v) gradient of acrylamide (Invitrogen) 0.1 μg of DdMCU-NTD (in the absence and presence of 0.5% (w/v) SDS) was analyzed by Blue native PAGE gel. The electrophoresis was first performed at 120 V in Anode Buffer (25 mM imidazole, pH 7.0) and Cathode Buffer A (7.5 mM imidazole, 0.02% (w/v) Coomassie blue G-250, 50 mM Tricine, pH 7.0) for 1 hr before switching Cathode Buffer A to Cathode Buffer B (7.5 mM imidazole, 0.002% (w/v) Coomassie blue G-250, 50 mM Tricine, pH 7.0) for another 4 hr. The Molecular Weight Marker (Merck #26610) was used as a reference.

### Ca^2+^ titration experiment for DdMCU-NTD

NMR titration experiments were performed at 303 K on a 14-T Bruker spectrometer equipped with a cryogenic TXI probe, operating at 599.702 and 60.774 MHz for ^1^H and ^15^N, respectively. A 500-μL sample containing 0.2 mM ^15^N-labeled DdMCU-NTD in 75 mM NaCl, 20 mM HEPES pH 7.5. Two-dimensional ^1^H-^15^N TROSY-HSQC spectra (Pervushin et al., 1997) were recorded at Ca^2+^ concentrations of 0, 1, 2, 5, and 10 mM. The spectra were processed by NMRPipe (Delaglio et al., 1995) and analyzed by CcpNmr (Vranken et al., 2005) to track the chemical shift changes induced by Ca^2+^.

## Results

### Structure determination of the DdMCU-NTD

For structural investigation of the NTD of DdMCU, we made five constructs, all lacking the predicted mitochondrial targeting sequence (residues 1-28), based on secondary structure and signal peptide prediction. These constructs include residues 29-126, 29-164, 29-164 with 88-106 removed (29-164/Δ88-106), 29-164/Δ92-106, 29-164/Δ93-106 with GGSGG added to the C-terminus (29-164/Δ93-106GGSGG) (Fig. S2). The synthetic dsDNAs of the constructs above were inserted into the pET21a vector between the *Nde*I and *Xho*I restriction enzyme sites for expression and thus all proteins have a 6×His tag at the N terminus. The proteins were expressed in *E. coli* and purified using nickel affinity, ion exchange, and size exclusion chromatography. The construct containing residues 29-126 showed the best homogeneity and stability and was used for crystallization.

We were able to crystallize DdMCU-NTD in a number of different conditions and in different forms. The best crystals were cube-shaped crystals formed in the precipitant solution containing 20% (w/v) PEG 4,000, 10% (v/v) 2-propanol, and 10 mM HEPES pH 7.5. These crystals diffracted to 1.7 Å with the P2_1_2_1_2_1_ space group. Since DdMCU-NTD has no sequence homology to any of the known structural domains in the database, we used the selenomethionine (SeMet) single-wavelength anomalous diffraction (SAD) method (Rice et al., 2000) for experimental phasing. DdMCU-NTD has only one methionine (Met) out of 98 residues. To avoid the potential problem of having too few Met sites for labeling, we introduced several methionine residues by mutagenesis, including L8M, L54M, L86M, L71M/L67M, L8M/L20M, L8M/L86M, and L8M/L20M/L86M. One of the mutants, L8M/L20M, successfully produced crystals with the P12_1_1 space group that diffracted to 2.2 Å. SAD data collected with the crystals were used to obtain the initial phase and to solve the SeMet mutant structure (Table 1). Four monomers with two types of interactions were found in an asymmetric unit (Fig. 1A) and all monomers were complete based on the electron density. One monomer was used as the search model to solve the wild-type structure by molecular replacement. The structure of the wild-type DdMCU-NTD was subsequently refined to 1.7 Å (Fig. 1B) with R_work_/R_free_ of 19.0/22.4% (Table 1). We focused our discussion on the P2_1_2_1_2_1_ structure because the protein was not mutated.

**Figure 1.**
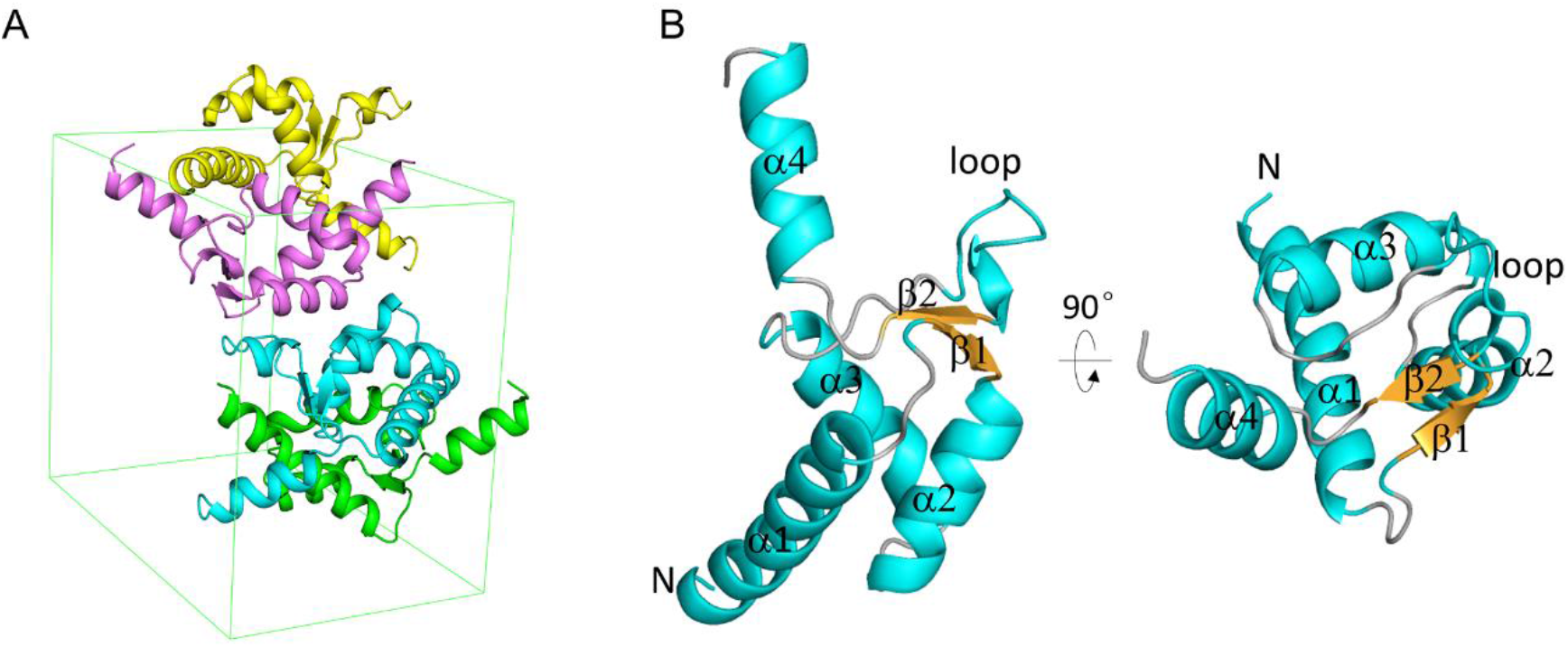
Overall structure of DdMCU-NTD. A. Ribbon representation of DdMCU-NTD with one pair of monomers colored green and cyan and one pair of monomers colored magenta and yellow. B. The crystal structure of DdMCU-NTD monomer is composed of four helices and two strands.

### DdMCU-NTD adopts a helix-rich fold

Each monomer in the DdMCU-NTD wild-type crystal contains four α-helices with two helix-loop-helix motifs and a short antiparallel β-sheet formed between segments of the two loops (Fig. 2A). This is distinctly different from all NTDs of the solved structures. Such structures, typicallized by the HsMCU, are largely composed of β-strands and adopt an ubiquitin-like fold (Fig. 2B). The NTDs of the four fungal homologues, MaMCU, CyMCU, NcMCU and NfMCU, show tertiary structural resemblance to HsMCU-NTD with RMSD 1.5-2.0 Å determined by Dali server (Fig. S3) in contrast to the low sequence identity/similarity (as low as 1.5~9.4%/2.4~13.6%) with HsMCU-NTD. Despite of the fungal origin, DdMCU-NTD shares neither sequence (Fig. S4) nor structure similarities with the structure-known fungal homologues, suggesting divergent evolution.

**Figure 2.**
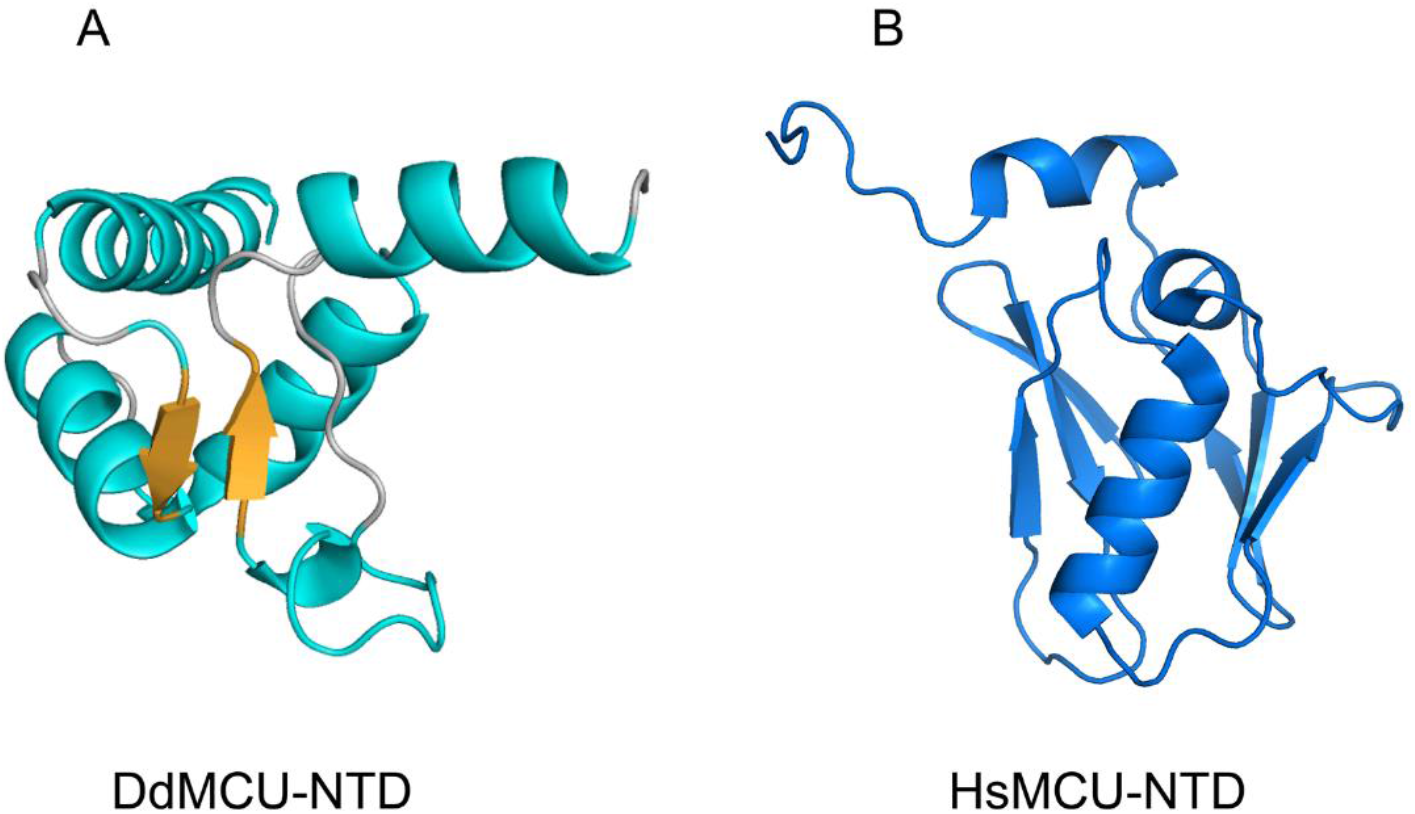
Crystal structure of DdMCU-NTD and comparison to the HsMCU-NTD structure. Ribbon representations of the crystal structures of (A) DdMCU-NTD (cyan) with two strands in yellow and (B) HsMCU-NTD (blue).

### DdMCU-NTD forms a stable homo-oligomer in solution

Consistent with various types of inter-monomer interactions found in the crystal structure, DdMCU-NTD was shown to form a higher-order oligomer in solution. The size exclusion chromatography (SEC) retention volume suggested a molecular weight (M.W.) of ~50 kDa, and a more accurate analysis with SEC-coupled multi-angle light scattering (SEC-MALS) suggested an average M.W. of 53.7 kDa (Fig. 3A), which is in close agreement with a 4- or 5-meric complex (M.W. of the theoretical 4- or 5-mer is 46.8 and 58.5 kDa, respectively). Higher-order oligomer of DdMCU-NTD is also supported by crosslinking and blue native polyacrylamide gel electrophoresis (BN-PAGE) results. Cross-linking experiments were further performed using the amine-reactive glutaraldehyde. Multiple bands with low migration rates were observed on the SDS-PAGE gel for the glutaraldehyde-treated sample, but not for the non-treated sample, suggesting a high-order oligomerization state (Fig. 3B). Next, we probed the oligomerization state of DdMCU-NTD using blue native polyacrylamide gel electrophoresis (BN-PAGE), which allows complexes to mostly maintain their stoichiometry. DdMCU-NTD migrated at a calibrated molecular weight of 63 kDa (Fig. 3C), which supported a higher-order oligomeric state of 5-/6-mer complex. Taken together, these results suggest that DdMCU-NTD forms an oligomer that is at least composed of four subunits and may exist as an equilibrium between pentamer and hexamer. By contrast, HsMCU-NTD forms mostly monomer and dimer in solution (Lee et al., 2016; Lee et al., 2015).

**Figure 3.**
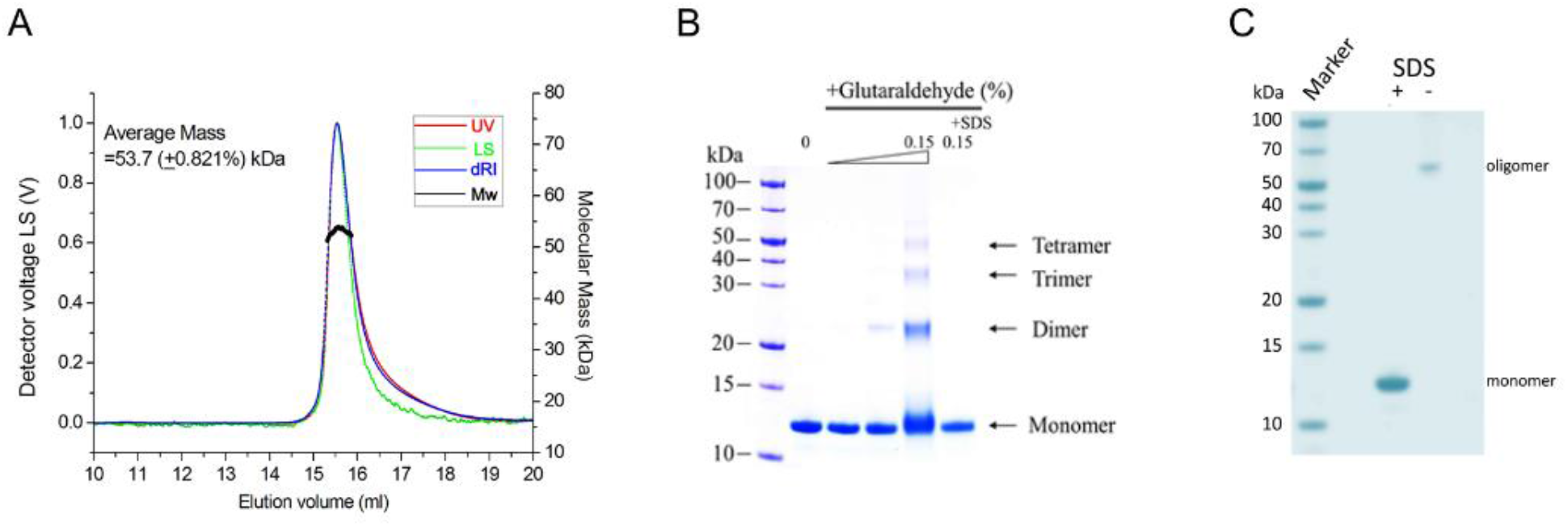
Oligomeric state of DdMCU-NTD. A. SEC-MALS analysis of DdMCU-NTD. The left and right axes represent the light scattering detector reading and molecular mass, respectively. The black curve represents the calculated molecular mass, and the average mass of the elution peak of DdMCU-NTD is 53.7 kDa. B. SDS-PAGE analysis of DdMCU-NTD after the crosslinking by glutaraldehyde. DdMCU-NTD was treated with increasing amount of glutaraldehyde for 2 min: 0, 0.0015%, 0.015% and 0.15% (v/v). For the comparison, the protein was treated with 1% SDS first and then with 0.15% gludealdehyde (the last lane). C. Blue Native PAGE gel of DdMCU-NTD indicates a high-order oligomer formation. The samples were loaded to blue native gradient gel with a 4-13% gradient of acrylamide in the presence and absence of 0.5% SDS.

### Key interactions driving NTD oligomerization in solution

A dimer of dimer DdMCU-NTD assembly was observed in the crystal unit that displays two types of intermolecular associations, antiparallel and parallel (Fig. 4, Fig. S5). To gain more insights into the interactions that mediate oligomerization in solution, we examined the two types of interfaces (Fig. S5) in the crystal structures by mutagenesis. The PISA program identified an antiparallel interface mediated by the intermolecular interactions including Gln38-Ile88, Gln38-Tyr111, Lys43-Glu115, Gln45-Ser85, and Gln45-Ser87 in the P2_1_2_1_2_1_ wildtype crystal structure. For example, the hydrogen bonds were formed between sidechain Hε of Gln38 from one monomer with Ile88 backbone carbonyl and Tyr111 hydroxyl from an adjacent monomer (Fig. 4A). Other intermolecular hydrogen bonds were formed between Gln45 and Ser85 and between Gln45 and Ser87 (Fig. S5). We introduced the single mutation Q38A and the double mutation S85A/S87A and found by size exclusion chromatography that neither disrupted DdMCU-NTD oligomerization in solution (Fig. 4B). Considering MCUs are parallel oligomers, the antiparallel packing is probably an artifact of crystallization.

**Figure 4.**
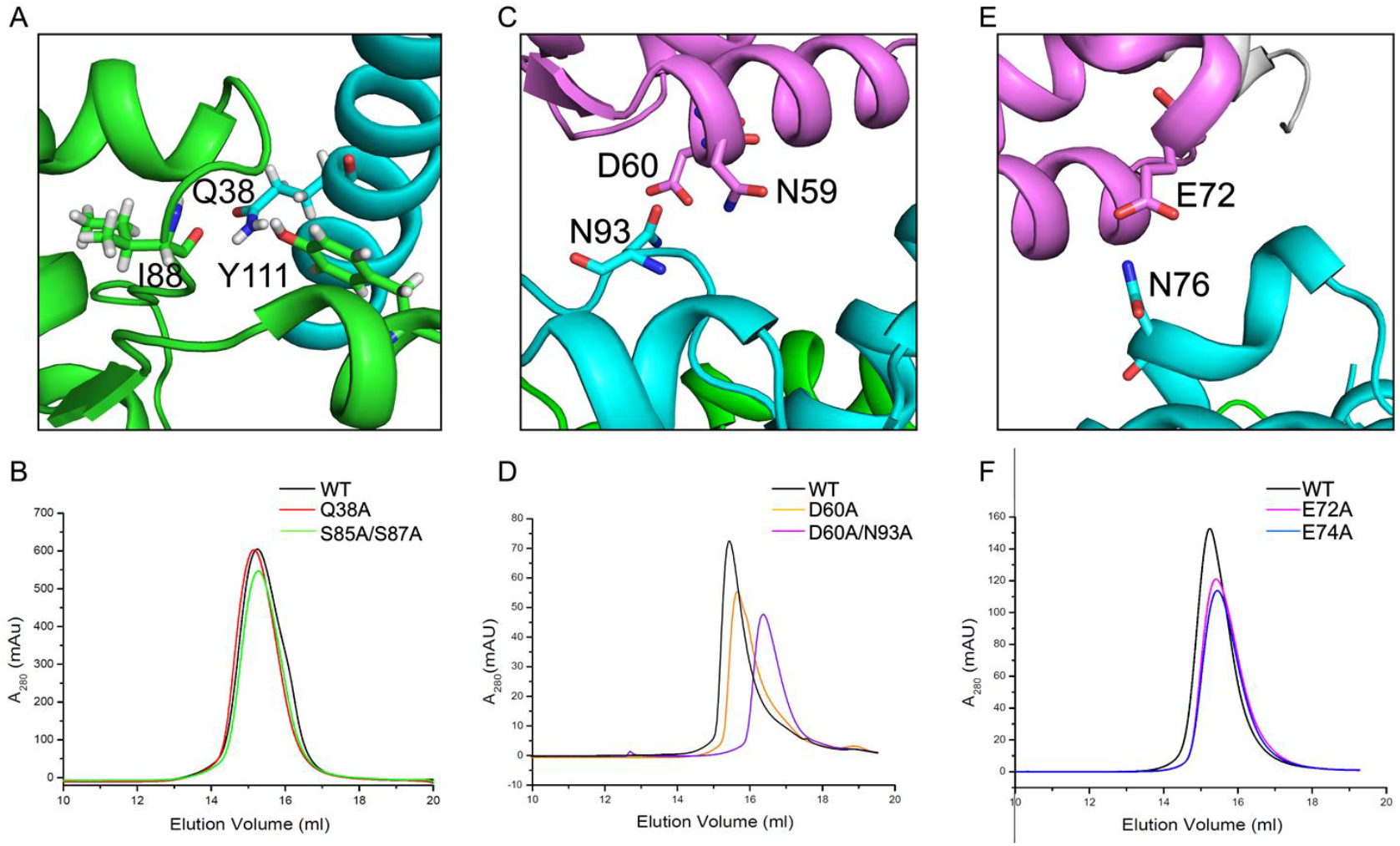
Key inter-molecular interactions for DdMCU-NTD oligomer formation. A. A detailed view of inter-molecular hydrogen bonds formed by Q38 with I88 and Y111. The two monomers are shown in green and cyan, respectively. B. Gel-filtration results of DdMCU-NTD wildtype and mutants. Elution peaks of DdMCU-NTD WT (black), Q38A (red) and S85A/S87A (green) from HiLoad16/60 Superdex200 column (GE Healthcare) in 75 mM NaCl, and 5 mM DTT, 20 mM HEPES pH 7.5. C. A detailed view of the inter-molecular N59/D60-N93 interactions. The oxygen and nitrogen atoms are shown in red and blue, respectively. D. Gelfiltration results of DdMCU-NTD WT (black), D60A (orange) and D60A/N93A (purple). E. A detailed view of the inter-molecular E72-N76 interactions. The oxygen and nitrogen atoms are shown in red and blue, respectively. F. Gel-filtration results of DdMCU-NTD WT (black), E72A (magenta) and E74A (blue).

Apart from the antiparallel interface, a noncrystallographic, parallel interface between two monomers also exists and appears to involve intermolecular interactions between Asn59 sidechain amine and Asn93 carbonyl, and between Asp60 sidechain carbonyl and Asn93 amine (Fig. 4C, Fig. S5). In this case, single mutations (D60A, N93A) or double mutation (D60A/N93A) disrupted the NTD oligomer (Fig. 4D). The SEC-MALS analysis of the D60A and N93A mutants showed apparent M.W. of 33.8 and 25.7 kDa, respectively. The double mutant D60A/N93A showed the greatest size reduction to 19.0 kDa (Fig. S6, Table 2). Another two pairs of the interactions in the parallel interface were between Glu72 sidechain carbonyl and Asn76 amine (Fig. 4E) and between Lys66 sidechain amine and Glu74 carbonyl (Fig. S5). We found that substituting the acidic sidechains of Glu72 and Glu74 also altered the oligomerization pattern (Fig. 4F), i.e., single mutations E72A and E74A reduced the apparent M.W. of the solution complex to 42.5 kDa and 39.8 kDa, respectively (Fig. S6, Table 2). All the mutated proteins were eluted as a single major peak on a Superdex 200 10/300 column with relatively broad peak width, except for the D60A. And the apparent molecular weights of the mutants ranged from ~19 kDa to ~42.5 kDa, equaling to 1.6 to 3.6 times the theoretical 11.7-kDa monomer mass, suggesting the existence of monomer-dimer, dimer-trimer, and trimer-tetramer equilibrium in fast exchange, respectively. Notably, mutating E71, which is spatially close to E72 and E74, did not cause any significant decrease in size, suggesting that the observed interface sites containing E72 and E74 are specific (Fig. S6). These results suggested that the parallel arrangement observed in the crystal structure mediated NTD oligomerization in solution.

**Table 2.** Molecular weights estimated by SEC-MALS (Related to Fig. S7)

| **protein** | **MW (kDa)<sup>a</sup>** | **SE** | **Mn (Da)<sup>b</sup>** | **MW/Mn<sup>c</sup>** |
| --- | --- | --- | --- | --- |
| WT | 53.7 | 0.8 | 11762 | 4.7 |
| WT+calcium | 20.2 | 0.1 | 11762 | 1.7 |
| Q38A | 58.7 | 1.4 | 11705 | 5.0 |
| S85A/S87A | 53.7 | 2.3 | 11730 | 4.6 |
| D60A | 33.8 | 0.9 | 11718 | 2.9 |
| N93A | 25.7 | 1.9 | 11719 | 2.2 |
| D60A/N93A | 19.0 | 2.7 | 11675 | 1.6 |
| E71A | 52.2 | 9.6 | 11704 | 4.5 |
| E72A | 42.5 | 1.4 | 11704 | 3.6 |
| E74A | 39.8 | 3.3 | 11704 | 3.4 |

<sup>a</sup>Average molecular mass (MW) of WT and mutants from the SEC-MALS data.
<sup>b</sup>Average molecular weight (Mn) of WT and mutants calculated based on the protein sequences.
<sup>c</sup>Ratio of the experimental molecular mass from SEC-MALS to the calculated molecular weight.
SE, standard error.

### Calcium and acidic residue mutations disrupt DdMCU-NTD oligomer in solution

A previous study of HsMCU-NTD showed that Ca^2+^ or Mg^2+^ could destabilize HsMCU-NTD oligomerization and shift the self-association equilibrium to monomer (Lee et al., 2016). We found similar Ca^2+^ response for DdMCU-NTD. In the presence of 50 mM Ca^2+^, the DdMCU-NTD oligomers reduced to an apparent M.W. of 20.2 kDa in MALS (Fig. 5A) and 22 kDa in DLS analysis (Fig. 5B). A control experiment using K^+^ showed no obvious effect on DdMCU-NTD oligomerization (Fig. S7), suggesting that the Ca^2+^-induced dissociation of the DdMCU-NTD oligomers is specific to divalent cations.

**Figure 5.**
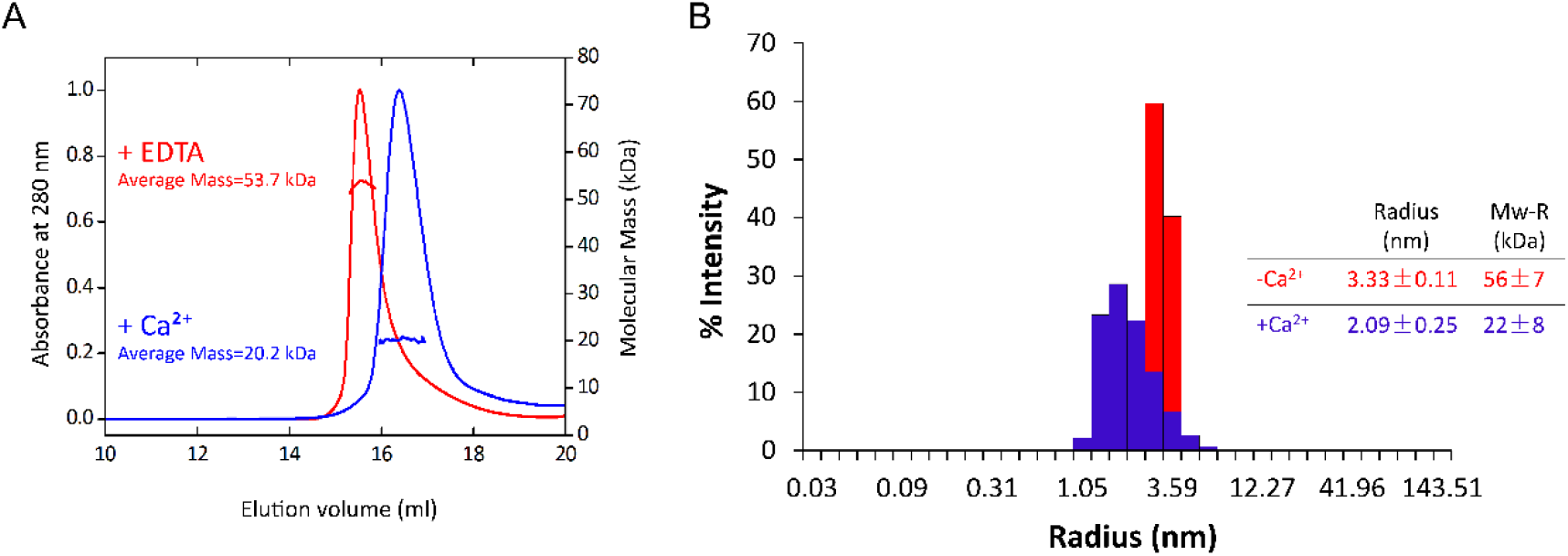
Modulation of DdMCU-NTD oligomerization by calcium. A. Gel-filtration analysis of the DdMCU-NTD in the absence (red) and presence (blue) of 50 mM Ca^2+^ from Superdex 200 10/300 GL column (GE Healthcare). B. DLS intensity particle size distribution of DdMCU-NTD in the absence (red) or presence (blue) of 50 mM Ca^2+^.

## Conclusions

As indicated from the sequence alignment, the DdMCU has higher sequence similarity in NTD-less (127-275) domain than the NTD (Fig. S4) with other homologues. The transmembrane pore formed by the TM helices contains the Ca^2+^ selectivity motif (DxxE). The conserved pore architecture reflects the ability of MCUs to mediate Ca^2+^ flux in a selective manner. Here we found that one major difference is the conformation of the NTD, whereas HsMCU-NTD adopts a β-grasp-like fold with 22.7% β strands, DdMCU-NTD has a helix-rich structure with 55.7%α helices. In addition to the crystal structure of DdMCU-NTD being different from that of HsMCU-NTD, the oligomerization of DdMCU-NTD in solution is also different from what was observed in HsMCU-NTD. Whilest the crystal structure showed both dimeric and tetrameric assembly, the DdMCU-NTD in solution showed higher-order oligomeric states when cross-linked. Native PAGE and SEC results also showed that DdMCU-NTD formed a 5- or 6-meric state. Because of the relatively low-molecular weight of DdMCU-NTD, and the modest accuracy of the abovementioned methods in probing oligomerization states, it is currently difficult to define the exact oligomeric state. It is also possible that the full-length protein will have different oligomerization states from the 5/6-meric states interpreted by our data using the truncated NTD. Although the oligomerization state of DdMCU-NTD is not definitive, it is clear that DdMCU-NTD can self-assemble into a high-order oligomer in solution, while HsMCU-NTD did not show evidence of a stable oligomeric assembly; it instead exhibited monomer-dimer equilibrium in fast exchange (Lee et al., 2016). We have independently confirmed by SEC and DLS that HsMCU-NTD in solution is predominantly dimeric (Fig. S8). Based on these results, we conclude that DdMCU-NTD has a different self-association state from HsMCU-NTD.

The recent published structures of the whole fungal MCUs show that MCUs form a well-packed tetrameric ring, while the NTDs are in a dimer of dimer assembly with two-fold symmetry. Two interfaces have been identified in these NTDs. The interfaces are stabilized by similar inter-subunit interactions including hydrogen bonding between polar side chains and a conserved salt bridge between a Lys side chain atoms of one subunit and an Asp side chain atoms of a neighbouring subunit (Fig. S9). Such polar interactions are also observed in the crystal structures of the HsMCU-NTD. Apart from the interfaces within one MCU channel, extensive dimerization interactions at the NTD were observed in the cryo-EM structure of MCU-EMRE complex that multiple sets of salt bridges and hydrogen bonds forms the dimerization interface between two MCU tetramers (Wang et al., 2019). In DdMCU-NTD crystal structures, three pairs of parallel polar interactions are also identified, including N59/D60-N93, K66-D74, E72-N76, which play an important role in the formation and stabilization of the DdMCU-NTD oligomers. Disruption of either pair of the polar interactions by point mutations dramatically decreased the size of DdMCU-NTD. Unlike the parallel polar interactions, the antiparallel interfaces observed in DdMCU-NTD did not appear to mediate the oligomerization within one DdMCU channel. Considering the unusual NTD assembly observed between two human MCU-EMRE complexes, it is possible that antiparallel interactions happen at the dimerization interface of two DdMCU channels. Thus instead of the sequence and structure diversities, the conserved oligomeric polar interface of the NTDs implies that the NTDs may play a role in maintaining the architectural integrity of the uniporter.

According to the discovery of Lee et al., the disruption of the MCU-regulating acidic patch (MRAP) of HsMCU-NTD by Ca^2+^ or Mg^2+^ could weaken the oligomer assembly of the NTD and promote the NTD monomerization (Lee et al., 2016). It is likely that Ca^2+^ exits from the membrane-proximal chamber after passing through the TM pore. In this case, the local concentration of Ca^2+^ in the chamber is estimated at the millimolar level. The observation led to a model in which the NTD of HsMCU plays an important Ca^2+^-dependent regulatory role. We also found that Ca^2+^ has the effect of shifting the self-association equilibrium of DdMCU-NTD in solution from 5-/6-mer to 1-/2-mer. According to NMR titration experiment (Fig. S10), Ca^2+^ binding did not induce major conformational change in the NTD monomer, which is consistent with the lack of Ca^2+^ binding in DdMCU-NTD using ITC (data not shown). A likely explanation of the Ca^2+^-induced oligomer dissociation is that Ca^2+^ can directly interact with acidic residues involved in oligomerization (e.g., D60, E72 and E74) (Fig. S6) and thus break the key electrostatic interactions that hold the DdMCU-NTD oligomer together. Similar Ca^2+^-induced NTD dissociation has been observed for DdMCU and HsMCU despite the large structural differences in the NTD, suggesting that the NTD may play a regulatory role in the Ca^2+^ conductance of MCU.

In conclusion, we have shown that the NTD of DdMCU adopts a fold that is completely different from the NTD of other MCU homologues. We have also shown that the DdMCU-NTD is capable of forming oligomers in solution and such oligomerization appears to be mediated by intermolecular electrostatic interactions involving several acidic residues. The binding of calcium to these acidic residues can disrupt the interactions and thus destabilize the DdMCU-NTD oligomers. Although the functional role of the NTD in DdMCU remains to be investigated, our results provides structural insights of the NTD in DdMCU channel assembly and the basis of calcium permeation.

## Accession numbers

Coordinates for the crystal forms have been deposited with the Protein Data Bank under accession codes of 5Z2H for the wild-type and 5Z2I for the SeMet labeled mutant.

## Financial supports

This work was supported by grants from a National key R&D Program of China (2017YFA0504804 to B. O.) and National Natural Science Foundation of China (U1732125 to B. O.), NIH Grant HL130143 (to J. J. C.), National Natural Science Foundation of China (U1632127 to D. L.).

## Acknowledgement

We thank the staffs from BL18U beamline of Shanghai Synchrotron Radiation Facility and the staff members of the Large-scale Protein Preparation System, the Mass Spectrometry System and the NMR facility at the National Facility for Protein Science in Shanghai (NFPS), Zhangjiang Lab, China for their instrument support and technical assistance in data collection and analysis.

## Author contributions

B. O., J. J. C., C. C., T. C. conceived of the study; C. C. identified the initial crystallization condition; Y. Y., M. W. and M. L. prepared samples; Y.Y. and M. W. performed crystallization and data collection under D. L.’s supervision; L. W., M. W. and J. W. analyzed the data and determined the structure; D. L., J. J. C. and B. O. wrote the paper and all authors contributed to editing of the manuscript.

## Abbreviations

BN-PAGE: blue native polyacrylamide gel electrophoresis
Ce: *Caenorhabditis elegans*
CHES: 2-(cyclohexylamino)ethanesulfonic acid
CMC: critical micelle concentration
Dd: *Dictyostelium discoideum*
Fg: *Fusarium graminearum*
Ma: *Metarhizium acridum*
Nc: *Neurospora crassa*
Cy: *Cyphellophora europaea*
Nf: *Neosartorya fischeri*
DLS: dynamic light scattering
DTT: dithiothreitol
EDTA: ethylenediaminetetraacetic acid
EMRE: essential MCU regulator
HEPES: (4-(2-hydroxyethyl)-1-piperazineethanesulfonic acid
Hs: *Homo sapiens*
IPTG: isopropyl-1-thiogalactopyranoside
LB: Luria-Bertani
MALS: multi-angle light scattering
MCU: mitochondrial calcium uniporter
MES: 2-(N-morpholino)ethanesulfonic acid
MICU: mitochondrial calcium uptake protein
MRAP: MCU-regulating acidic patch
M.W.: molecular weight
NMR: nuclear magnetic resonance
NTD: N-terminal domain
OD_600_: optical density at 600 nm
PMSF: phenylmethylsulfonyl fluoride
SAD: single-wavelength anomalous diffraction
SEC: size exclusion chromatography
SeMet: selenomethionine
SDS: sodium dodecyl sulfate
SSRF: Shanghai Synchrotron Radiation Facility
TM: transmembrane

