## Supplemental figures for "Structural characterization of the N-terminal domain of the *Dictyostelium discoideum* mitochondrial calcium uniporter"

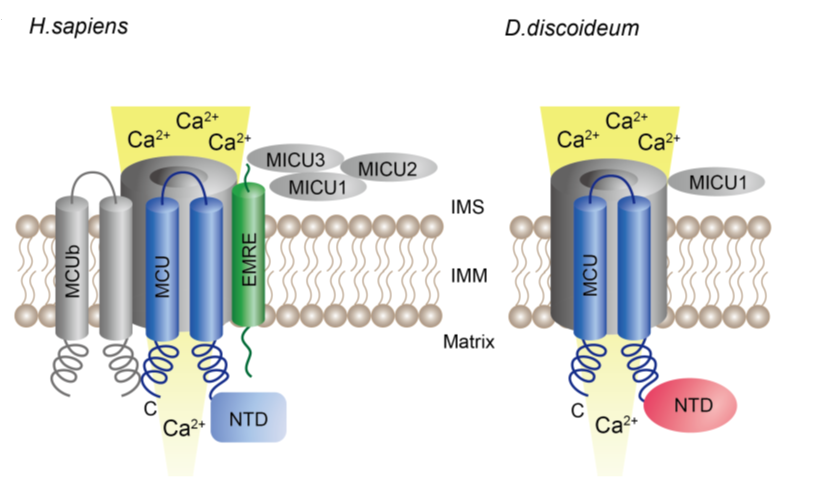
**Figure S1.** The components of the mitochondrial calcium uniporter complexes in *Homo sapiens* and *Dictyostelium discoideum*. The uniporter components in *Homo sapiens* consist of mitochondrial calcium uniporter protein (MCU), MCU regulatory subunit b (MCUb) and essential MCU regulator (EMRE), together with the intermembrane space (IMS) proteins mitochondrial calcium uptake protein 1 (MICU1), MICU2 and probably MICU3. The *Dictyostelium discoideum* uniplex proteins only contain MCU homologue and a putative MICU1 homologue. The major difference for the pore-forming component MCU in *Homo sapiens* and *Dictyostelium discoideum* lies in the N-terminal domain (NTD).

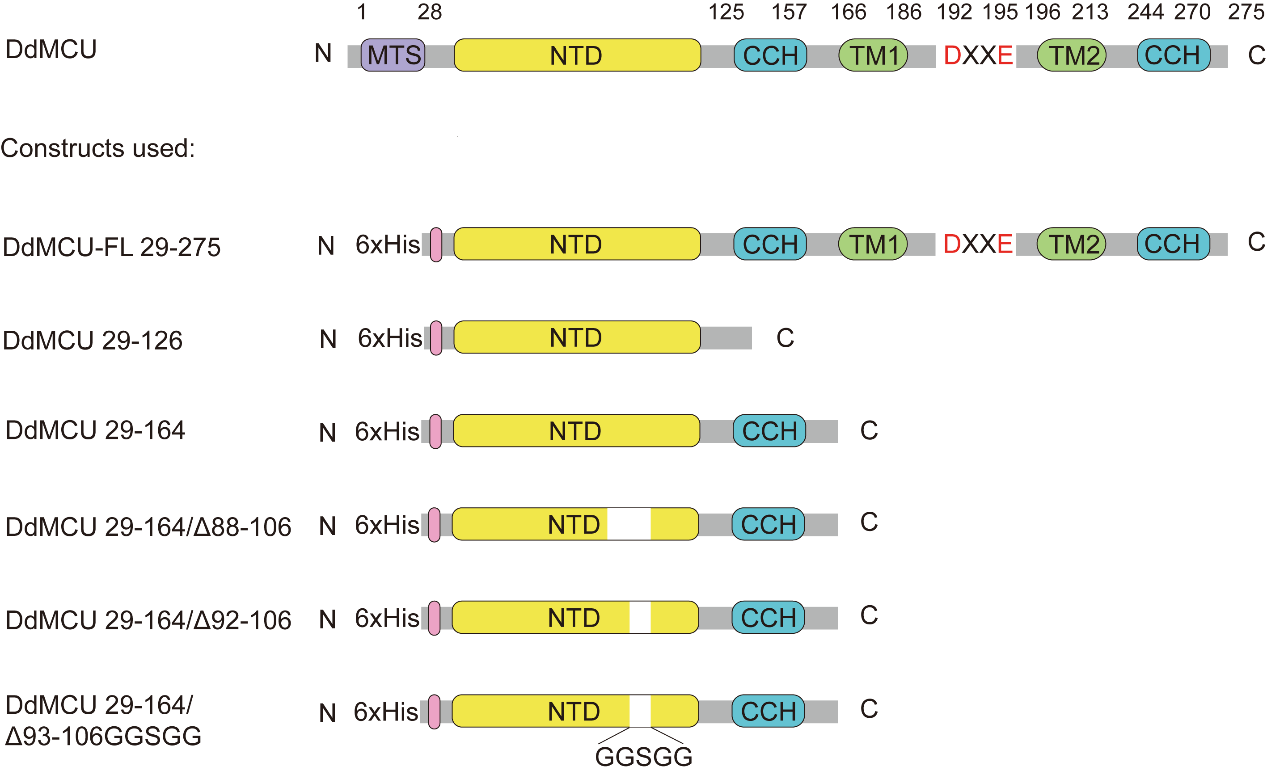
**Figure S2.** A linear overview of the domain organization for the original sequence of DdMCU and the constructs used in this paper. For each protein construct, different colors represent different domain: the predicted mitochondrial targeting signal (MTS) (purple), 6×His tag (pink), N-terminal domain (NTD) (yellow), transmembrane domain (TM) (green), the conserved DXXE motif, and coiled-coil helix (CCH) (blue).

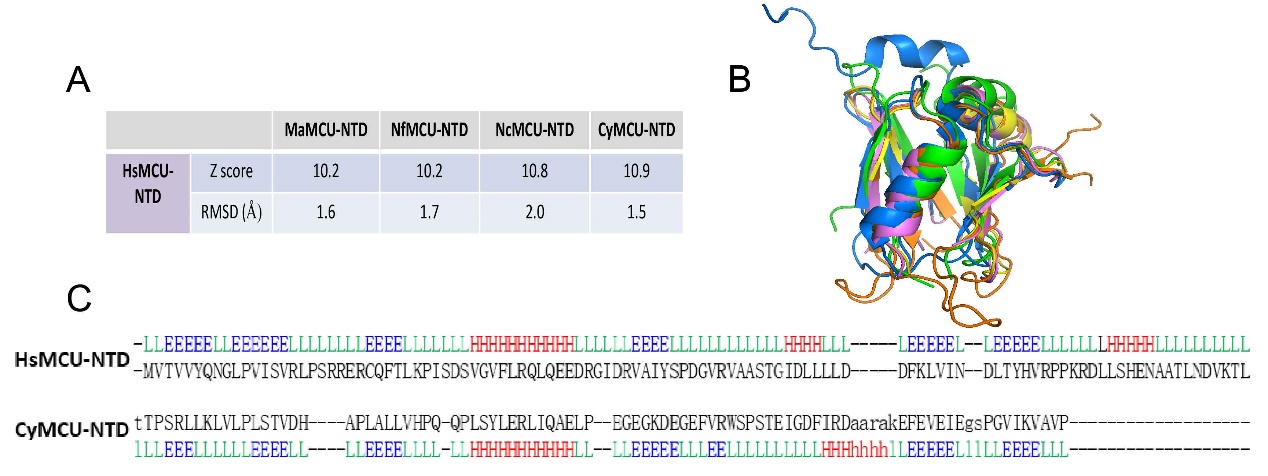
**Figure S3.** Structural alignment of MCU with different species homologues. Dali server was used to generate the structure alignment. (A) The z-score and RMSD values of different fungal homologues comparing to HsMCU-NTD. (B) Superimposing protein structures of HsMCU-NTD (blue) with MaMCU-NTD (PDB code: 6C5R) (yellow), CyMCU-NTD (PDB code: 6DNF) (violet), NcMCU-NTD (PDB code: 6DT0) (green), NfMCU-NTD (PDB code: 6D7W) (orange) (C) The aligned secondary structure between HsMCU-NTD and CyMCU-NTD (H/h: helix, E/e: strand, L/l: coil). Uppercase means structurally equivalent positions with CyMCU-NTD. Lowercase means insertions relative to CyMCU-NTD.

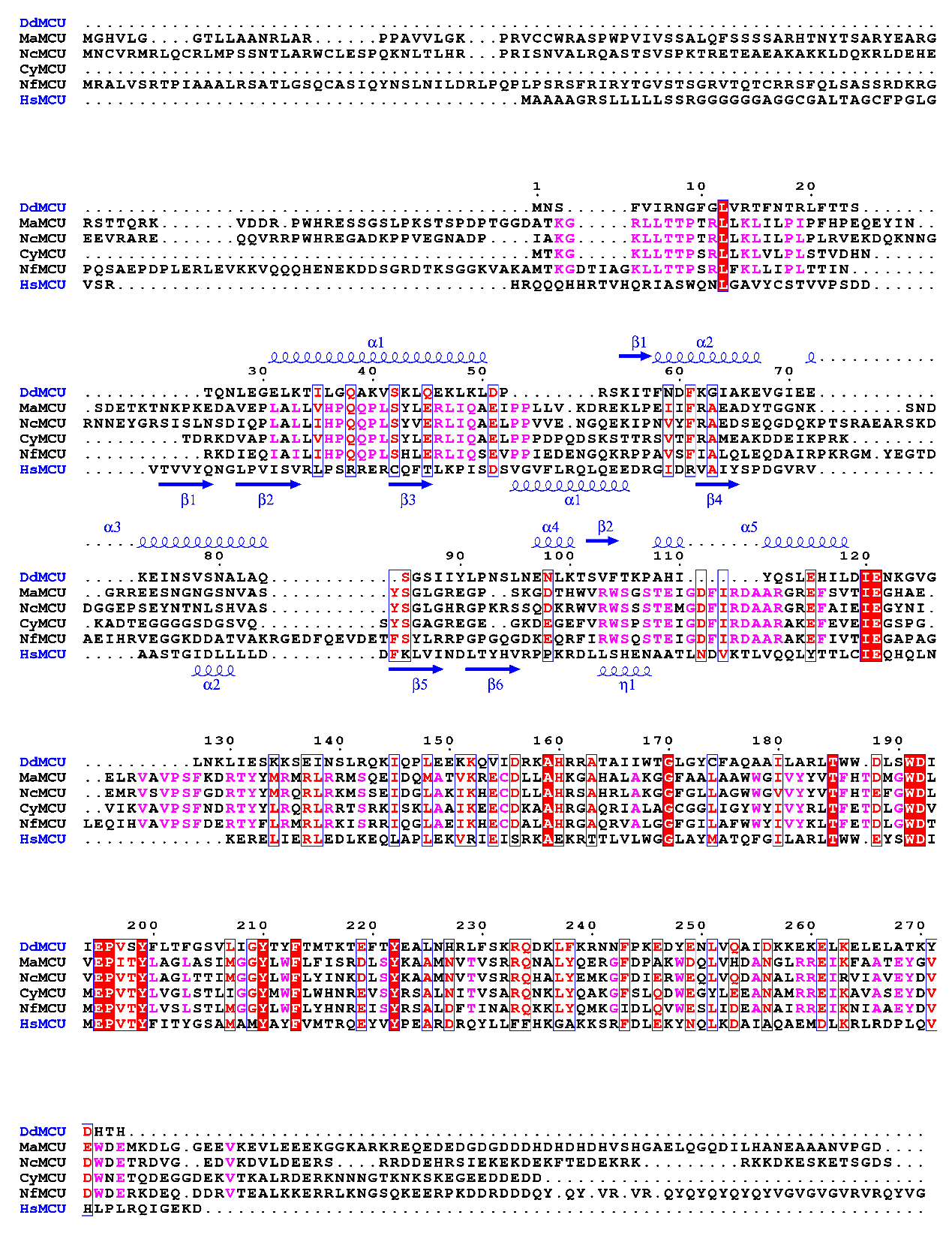
**Figure S4.** Sequence alignment of different MCU homologues. The comparison of the sequence HsMCU with the fungal homologues and the comparison of the secondary structures of DdMCU-NTD and HsMCU-NTD using ClustalW and ESPript (Robert and Gouet, 2014) programs.

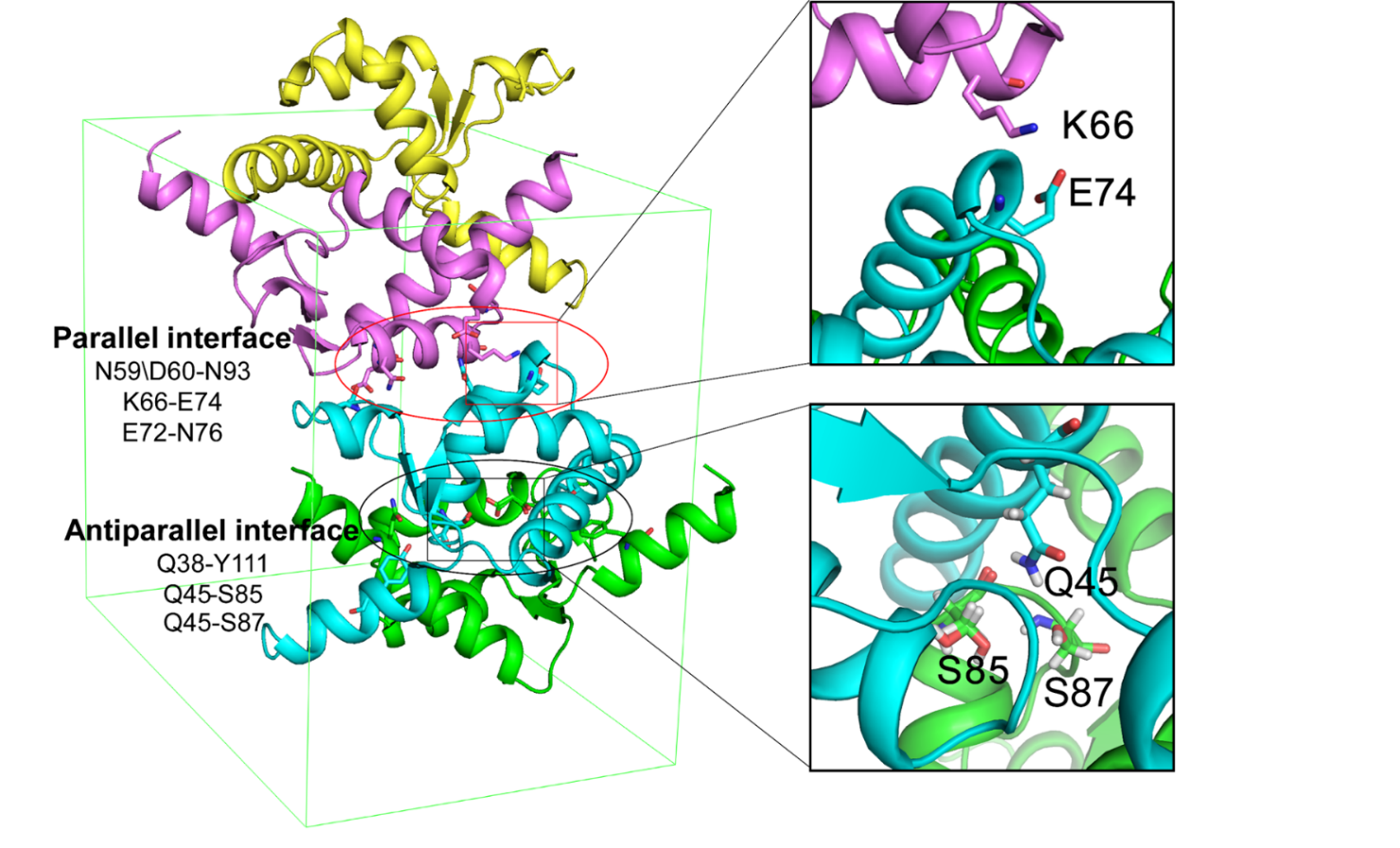
**Figure S5.** The packing interfaces of DdMCU-NTD in the unit cell. Two interfaces are identified, including parallel interface (red) containing three pairs of inter-molecular interactions formed by N59/D60-N93, K66-E74, E72-N76 and antiparallel interface (black) containing two pairs of inter-molecular hydrogen bonds formed by Q38-I88/Y111, and Q45-S85/S87. Detailed views of the pairs of K66-E74 and Q45-S85/S87 are shown on the right side.

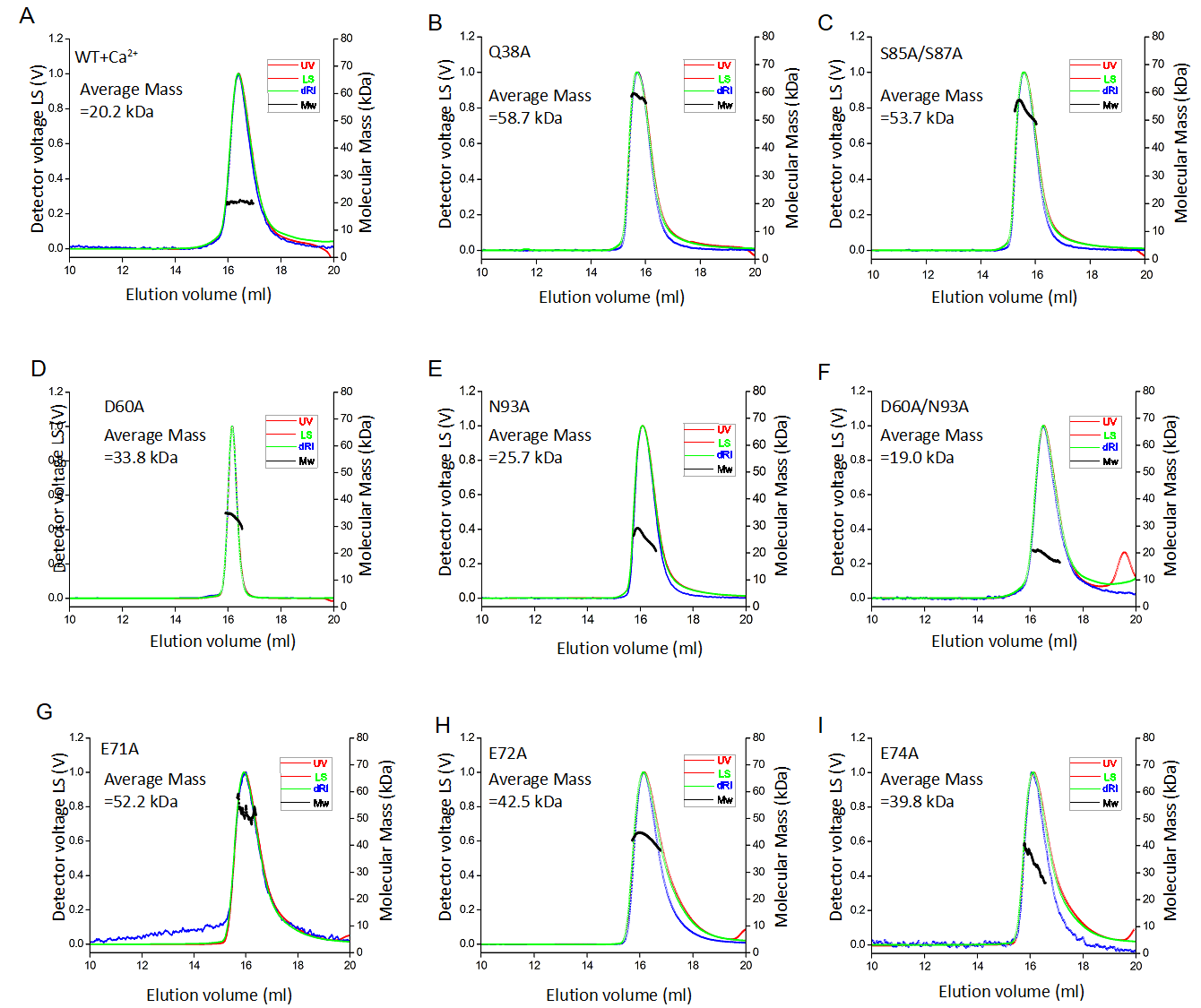
**Figure S6.** SEC-MALS analysis of different mutants. Chromatograms show the readings from the light scattering at 90° (green), refractive index (blue), and UV (red) detectors. The left and right axes represent the light scattering detector reading and molecular mass, respectively. The black curve represents the calculated molecular mass, and the average mass of the elution peaks of mutants Q38A, S85A/S87A, D60A, N93A, D60A/N93A, E71A, E72A, and E74A are 58.7 kDa, 53.7 kDa, 33.8 kDa, 25.7 kDa, 19.0 kDa, 52.2 kDa, 42.5 kDa and 39.8 kDa, respectively. Calcium addition to WT shifts the average molecular mass to 20.2 kDa.

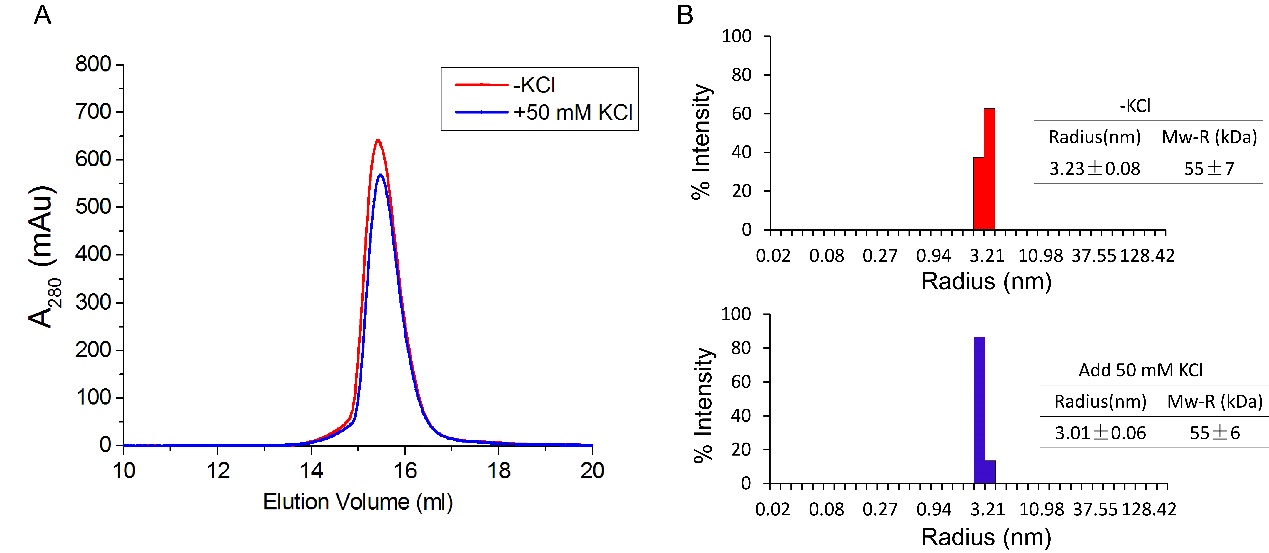
**Figure S7.** Potassium effects on DdMCU-NTD oligomerization. A. Gel-filtration analysis of the DdMCU-NTD in the absence (red) and presence (blue) of 50 mM K^+^ (Superdex 200 10/300 GL column (GE Healthcare)). B. DLS intensity particle size distribution of DdMCU-NTD in the absence (red) or presence (blue) of 50 mM K^+^.

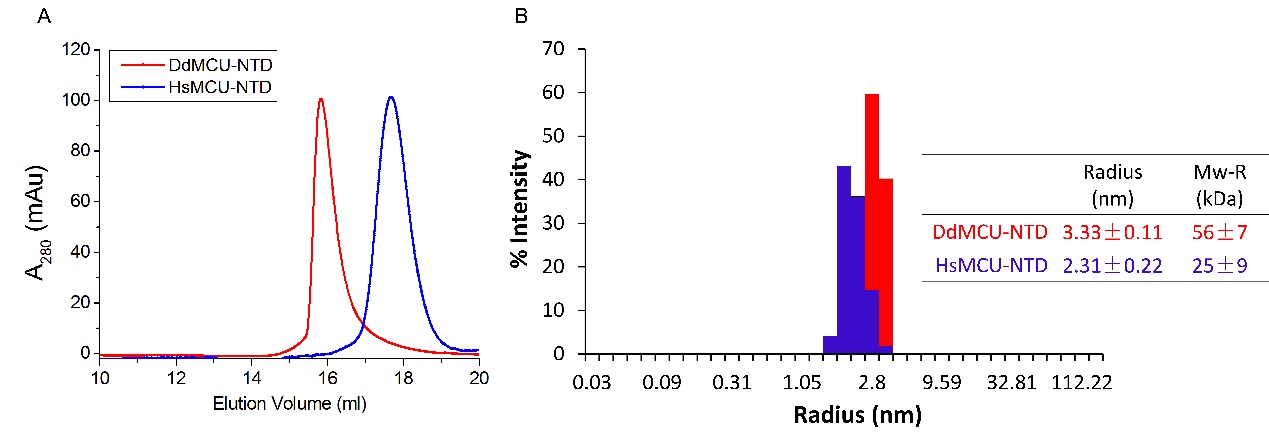

**Figure S8.** The oligomerization of DdMCU-NTD and HsMCU-NTD. A. Gel-filtration results of DdMCU-NTD and HsMCU-NTD. B. DLS results of DdMCU-NTD and HsMCU-NTD.

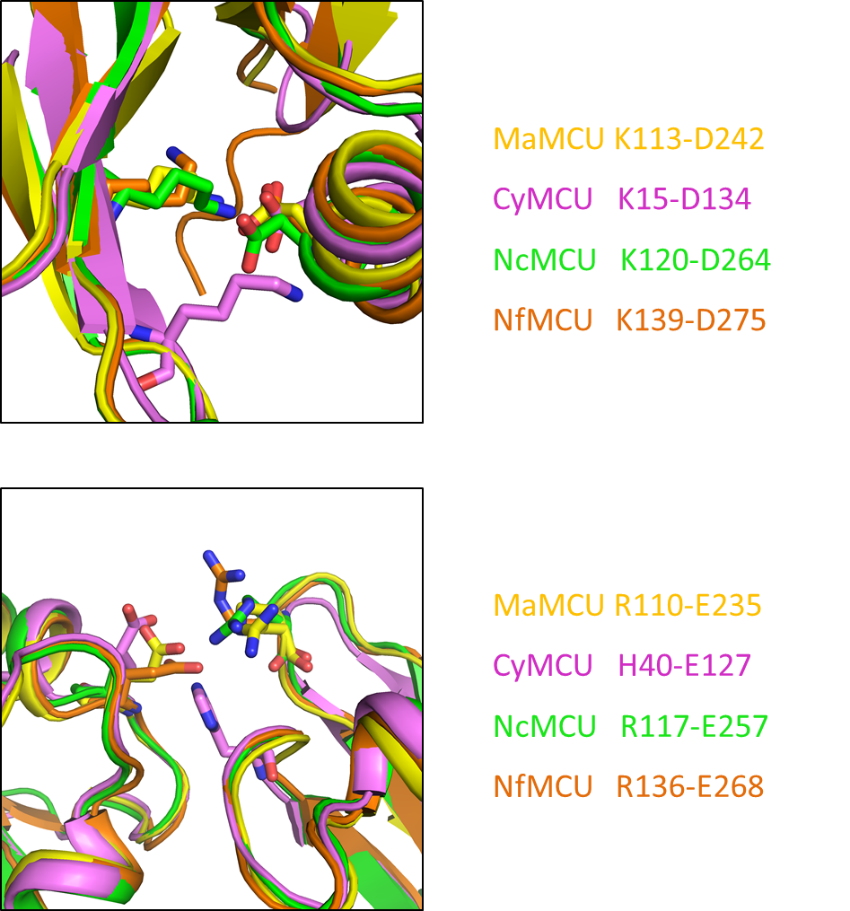
**Figure S9.** The conserved atomic interaction in the interface of two NTD subunits of the four fungal homologues. The salt bridge formed between an Asp and a Lys is identified in MaMCU-NTD (yellow), CyMCU-NTD (violet), NcMCU-NTD (green), NfMCU-NTD (orange).

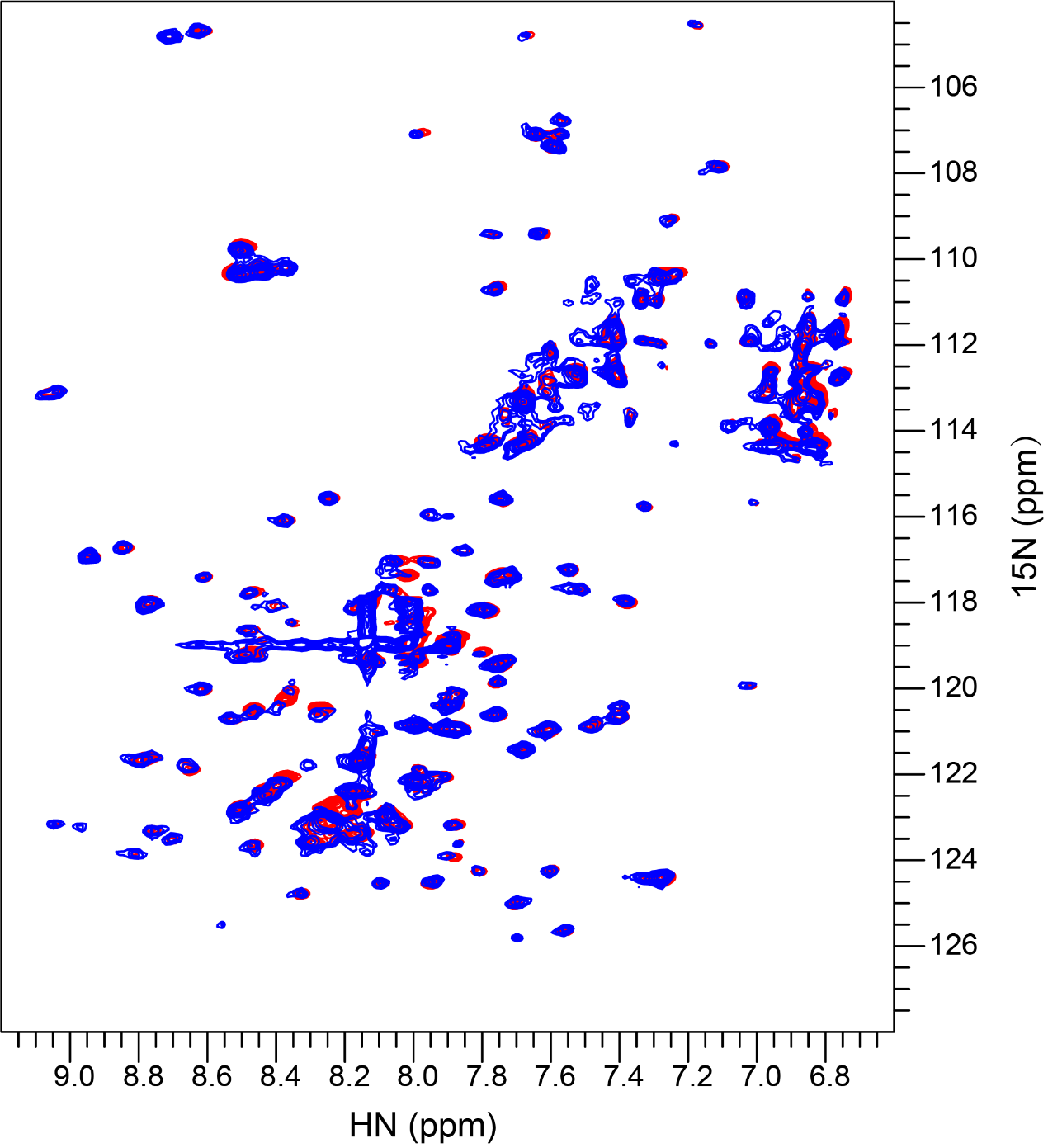
**Figure S10.** The calcium titration showing the chemical shift changes in the absence of calcium (red) and in the presence of 5 mM calcium (blue). The minor changes observed between the two spectra indicate that the overall structure of DdMCU-NTD doesn’t change upon the calcium addition.
